# Bulky lesions on the displaced strand stimulate DNA unwinding by dimeric UvrD-family helicases

**DOI:** 10.64898/2026.08.03.742518

**Authors:** Eric J. Tomko, Ankita Chadda, Eric A. Galburt

## Abstract

UvrD-family SF1A helicases play a variety of biological roles in DNA metabolism including replication, recombination, DNA repair, and conjugative plasmid transfer. The family is described by a subdomain architecture consisting of two RecA motor subdomains (1A and 2A) that each contain an auxiliary B-domain insertion (1B and 2B). Monomeric UvrD-family enzymes possess ATPase and 3’-5’ single-stranded DNA translocase activity but lack helicase activity. An enzyme dimer formed through a 2B-2B domain interface acts as a processive DNA helicase. A partial explanation for this observation, based on structural comparison between monomeric and dimeric DNA bound complexes, is that dimerization removes inhibitory contacts between the 2B domain and the double-stranded DNA, thus activating the helicase. However, biochemical observations reveal additional aspects of the dimeric mechanism that cannot solely be explained by movement of the 2B subdomain. Here, we present data showing that the *Mycobacterium tuberculosis* UvrD1 dimer interacts with the displaced strand of the duplex (*i*.*e*., the 5’-3’ strand in the direction of unwinding). In particular, bulky lesions on the displaced strand–including a thymine dimer–led to more efficient unwinding over a finite range of duplex lengths. These findings suggest a model for dimeric UvrD-family unwinding wherein one subunit makes intimate contacts with the displaced strand and that this interaction promotes the processive DNA unwinding unique to dimers.

## Introduction

Helicases are enzymes that couple NTP hydrolysis to nucleic acid strand separation (*i*.*e*., unwinding). They are involved in mediating processes such as replication, recombination, transcription, translation, RNA splicing, and DNA repair. These enzymes can be classified according to sequence and structural conservation into six superfamilies (SF) [1,2]. The SF1 superfamily is further subdivided depending on the direction of translocation on single-strand DNA. SF1A enzymes such as UvrD, UvrD1, Rep, PcrA, RecB and Srs2 exhibit 3’ to 5’ directionality while SF1B enzymes such as RecD, Pif1 and Dda move in a 5’ to 3’ direction. A subset of the SF1A enzymes belongs to the UvrD-family as determined by close homology with *Escherichia coli* UvrD DNA helicase [3].

UvrD-family enzymes use the free energy from ATP binding and hydrolysis to power processive 3’ to 5’ translocation along single-stranded DNA (ssDNA) [4–6]. The ATP binding site is formed between the two RecA-like motor subdomains (1A and 2A), and conformational changes coupled to the ATPase cycle are coupled to directional movement [7]. Single-stranded DNA translocation is itself functional and is implicated in the reorganization and removal of proteins from ssDNA during replication and recombination [8– 10]. Crucially, while monomeric UvrD-family enzymes are active ssDNA translocases, they must be activated to become processive helicases. This activation can occur via auxiliary protein partners (*e*.*g*., UvrD by the mismatch repair protein, MutL [11] and Rep by the replication restart protein, PriC [12]), force (*e*.*g*., in the context of single molecule experiments [13]), or — most fundamentally —by dimerization [14–19].

The dimerization interface responsible for helicase activation was recently identified between the 2B sub-domains of each subunit in *Mycobacterium tuberculosis* UvrD1 and *E. coli* (*Ec*) UvrD [19–21]. This sub-domain has been shown to exhibit rotational conformational flexibility around its linker connections to the 2A motor domain and relative to the rest of the protein as large as 160 degrees [20,22–26]. In the context of the *Mycobacterium tuberculosis* (*Mtb*) UvrD1 dimer bound to a ssDNA-dsDNA junction [20,21], each domain adopts unique rotational positions, leading to the removal of inhibitory 2B-dsDNA contacts present in DNA-bound monomer complexes. Consistent with this structural interpretation, the 2B domain position was previously implicated in the activation of *Ec*UvrD via dimerization [26] and by MutL interaction [11], as well as in the activation of Rep via PriC [12]. In addition, deletion of the Rep 2B sub-domain activates helicase activity in a monomer, albeit with slower kinetics and processivity [27].

However, while structural rearrangement of the 2B sub-domain is necessary for helicase activation, it cannot explain additional properties of dimer-catalyzed DNA unwinding. For example, the ATPase activity of Rep is enhanced upon dimerization, suggesting linkage between dimerization and ATP hydrolysis [28,29]. In addition, both subunits of UvrD and PcrA must be able to hydrolyze ATP [16,18,26], and the two ATPase sites within each subunit are functionally coupled [30]. Furthermore, tryptophan mutants of Rep that can dimerize, but show no helicase activity, are able to activate WT subunits through hetero-dimer formation [30]. One question that stems from these observations is whether individual subunits within a dimer perform distinct roles during unwinding–a possibility further promoted by the unique conformations of each subunit observed in cryoEM structures [20].

In fact, many mechanistic questions remain regarding UvrD-family dimeric helicase activity. Experiments suggest that the lead subunit must interact directly with a region of the duplex DNA [28,31], but the nature of this interaction is not known. It also remains to be determined whether the subunits change relative positions (*i*.*e*., leading and trailing) during processive translocation. Both inchworm models where the subunits do not exchange positions and rolling (or hand-over-hand) models have been considered [28,30,32,33].

Another unresolved mechanistic aspect is whether the helicase interacts with one or both DNA strands. For example, early work investigating the effects of a DNA-bound chemical adduct (CC-1065) on multi-round DNA unwinding showed that while SF1B helicase T4 Dda was only inhibited with the adduct on the loading strand, *E. coli* UvrD was also inhibited when the adduct was on the displaced strand [34]. Amongst possible mechnanisms, the possibility that UvrD interacts with both strands was discussed. Specific variations of such a model would include the dimer maintaining contact with both unwound strands (in a topology similar to the unrelated SF1A/SF1B RecBCD complex [35]) or an individual subunit interacting with the to-be-displaced strand in the context of the downstream duplex while the other maintains the canonical contacts with the 3’ to 5’ loading strand. We set out to formally test this entire class of mechanisms by performing DNA unwinding experiments with DNA containing bulky adducts within the displaced (5’ to 3’) strand. Our data provide strong evidence of an unappreciated interaction with the displaced strand and new hints as to the properties of UvrD dimers that facilitate their processive helicase activity.

## Results

### Bulky modifications in the backbone of the displaced strand activate unwinding

UvrD-family enzymes translocate on ssDNA in a 3’-5’ direction. Consequently, double-stranded (ds) DNA substrates used for unwinding studies use a 3’ ssDNA tail at one or both ends to allow for enzyme binding and subsequent translocation towards the duplex. We will refer to this strand as the loading strand and distinguish it from the displaced strand which sits 5’-3’ in the direction of unwinding. As described in the introduction, dimers of UvrD1 are required for helicase activity in the absence of other factors. As such, all experiments described here are performed under oxidative conditions where the covalent dimeric UvrD1 fraction is between 10-30% of the total enzyme concentration unless otherwise specified. Comparisons across different DNA substrates are always made with a constant fraction of dimer. Control experiments with 100% monomeric, prepared by inclusion of reducing agent, showed no DNA unwinding as previously reported [19]. Buffer conditions for unwinding experiments were 20 mM TRIS-HCl, pH 8.0 (25 °C), 75 mM NaCl, 20% (v/v) glycerol, 5 mM MgCl_2_, 1 mM ATP, and 5 μM ssDNA trap unless otherwise specified.

We constructed two DNA substrates consisting of either 18 or 40 base-pairs (bp) of duplex DNA and a 3’ tail consisting of 20 deoxythymidine nucleotides (dT_20_) (Figure 1A, black and blue respectively, see Supplemental Table 1 for a list of substrate sequences). The downstream ends of the loading and displaced strands were labeled with Cy5, and a black hole quencher (BHQ) respectively, allowing for duplex unwinding to be monitored via increases in Cy5 fluorescence [19]. The experimental fluorescence signal is converted into fraction unwound by comparing it to controls of both fully hybridized and fully single-stranded loading strand (Methods). Fractions unwound reported were taken after the signal had reached a plateau at 10 seconds.

**Figure 1:**
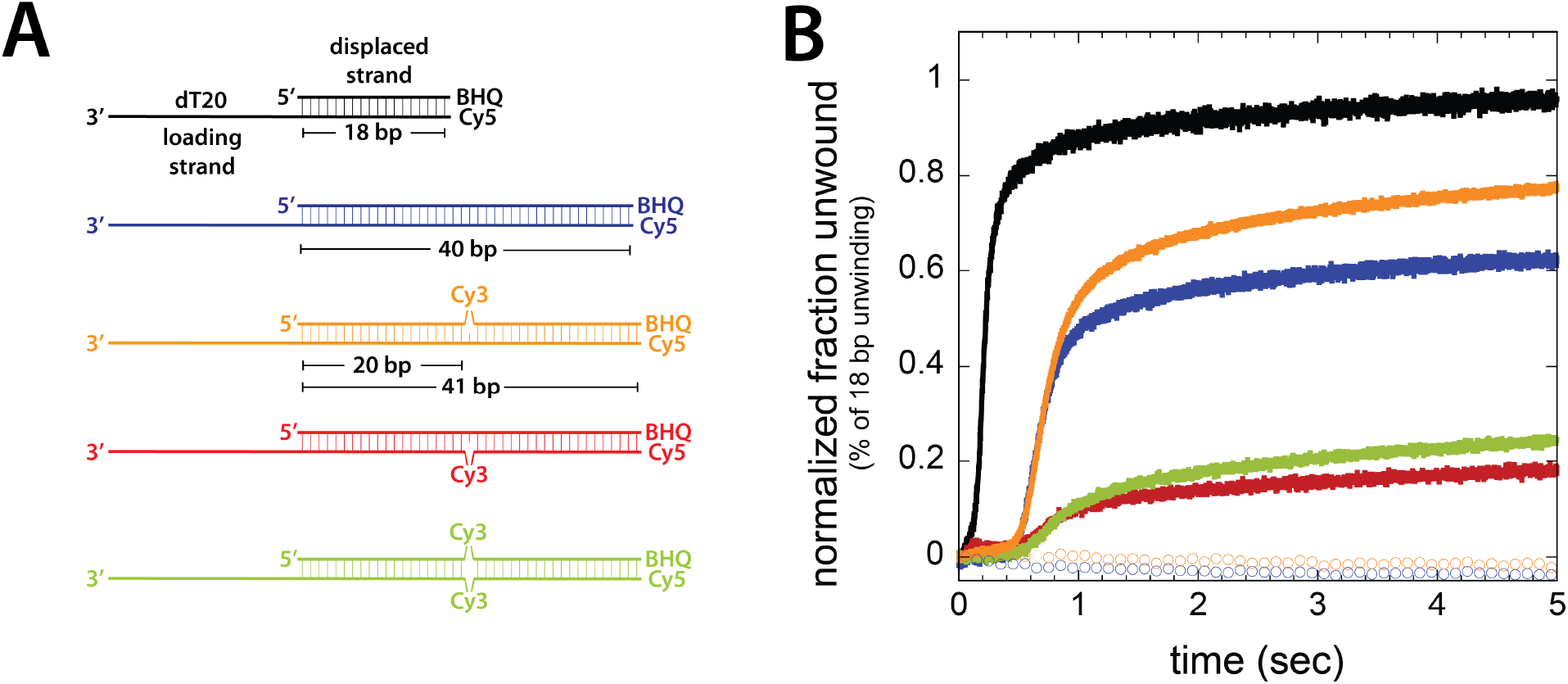
The effects on unwinding of various Cy3 modified substrates. **(A)** Different substrates with lengths and positions of Cy3 backbone modification indicated. Displaced strands are always drawn on top while tailed, loading strands on the bottom. **(B)** Stopped-flow DNA unwinding traces for each substrate are shown in solid lines of the corresponding colors while control reactions using monomeric UvrD1 (C451A) are shown in open circles.

Under single-round reaction conditions in the presence of 200 nM protein (monomer units), 2 nM DNA, and an excess of unlabeled ssDNA trap, 13.4 ± 0.4 % of the 18 bp substrate and 8.6 ± 1.0 % of the 40 bp substrate were unwound by UvrD1 (Supplemental Figure 1). The decreased unwinding on the longer template is expected due to the finite processivity (*i*.*e*., the probability of the enzyme to take a step forward as opposed to dissociating from the DNA). The absolute unwinding amplitude under these conditions is limited by the difficulty in generating 100% dimeric UvrD1 and the competition for DNA binding by inactive monomers. This is evident by the reduction in unwinding amplitude resulting from titrating in mutant UvrD1 (C451A) which is an obligate monomer (Supplemental Figure 2A,B) [19]. To compare relative amounts of unwinding between different substrates, we normalize the fraction DNA unwound to the 18 bp substrate amplitude (Figure 1).

We then examined a substrate similar to the 40 bp duplex, but with a Cy3 molecule placed in the backbone of the displaced strand in the middle of the duplex (Figure 1A, orange). Based on our initial hypothesis when beginning this work that UvrD1 does not interact with this strand during unwinding, our expectation was that this modification would have no effect on unwinding and previous observations using the CC-1065 adduct may have predicted an inhibition of unwinding [34]. Instead, we observed that the reaction was significantly stimulated, and that the fraction of unwound DNA increases by 23 ± 3% (Figure 1B, orange compared to blue)). Under monomeric conditions using UvrD1(C451A), neither of the substrates were unwound, consistent with the stimulatory effect being due specifically to the dimer (Figure 1B, open circles). In addition, while these experiments were performed under conditions of excess protein (200 nM UvrD1 and 2 nM DNA after mixing) to maximize the unwinding signal, experiments performed with excess DNA (10 nM UvrD1 and 100 nM DNA after mixing) displayed a similar enhancement of (26 ± 5%), demonstrating that the effect is only dependent on a single UvrD1 dimer on each substrate without contributions from larger oligomers or multiple UvrD1 dimers (Supplemental Figure 2C-E).

In contrast to the enhancement observed above, a substrate with Cy3 placed in the backbone of the loading strand inhibited unwinding and resulted in a decrease in the fraction unwound (Figure 1, red compared to blue). Substrates with Cy3 inserted into both strands also displayed reduced unwinding suggesting that the effect of the Cy3 in the loading strand is dominant (Figure 1, green compared to blue).

To ascertain whether these effects were specific to Cy3, we repeated the experiments using different modifications (Figure 2). Both a biotin linked to a dT nucleobase via a 12-carbon chain and a short backbone polyethylene glycol spacer (PEG 3) also resulted in activation (30 ± 10 % and 20 ± 8 % respectively, Figure 2A, diamonds and circles respectively). However, neither an abasic site nor a dU:dG mismatch led to stimulation of unwinding (Figure 2A, squares and triangles respectively).

**Figure 2:**
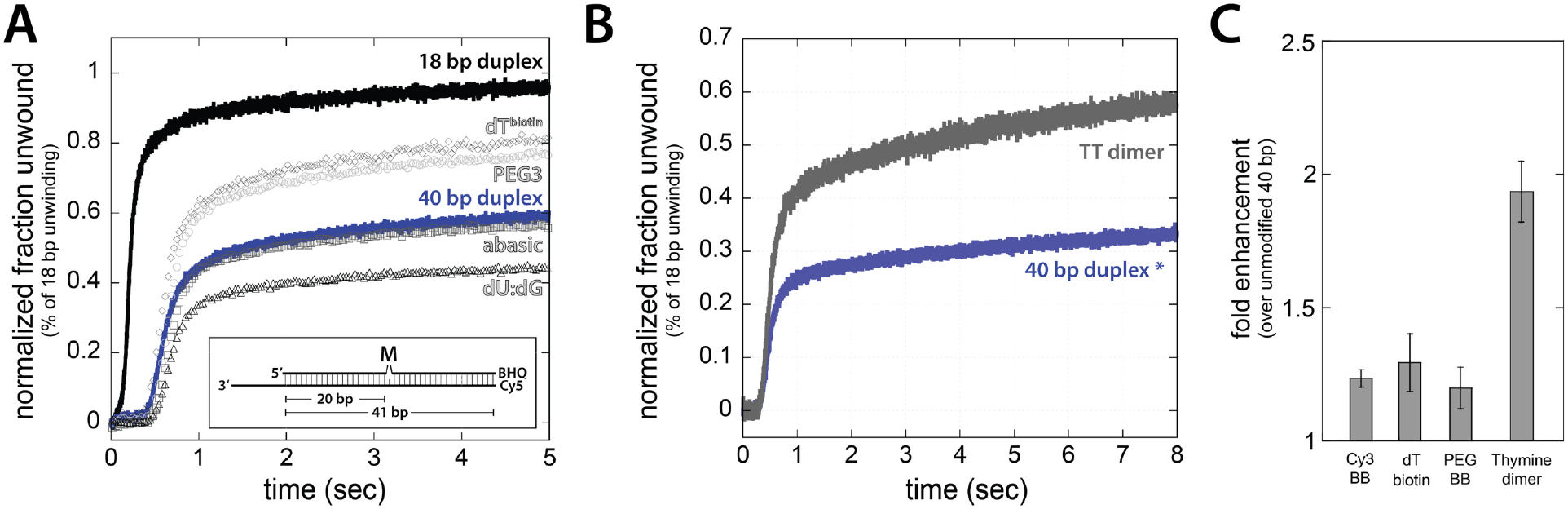
The effects on DNA unwinding of different internal modifications in the displaced strand. **(A)** Normalized unwinding traces using templates modified at position M with dT-biotin (diamonds), PEG3 (circles), abasic (squares), and dU:dG mismatch (triangles) compared to control substrates of 18 bp (black) and 40 bp (blue). See Supplemental Tables 2 and 3 for a list of sequences. **(B)** Fraction unwound as a function of time on the thymine dimer substrate (gray) compared to the control 40 bp (blue) substrate. **(C)** Fold-increase in unwinding for different modifications. BB indicates backbone modifications and the thymine dimer fold enhancement is shown corrected for the modified DNA fraction.

### Thymine dimers in the displaced strand dramatically enhance unwinding

Given the link between bulky lesions that distort double-stranded DNA structure and nucleotide excision repair (NER) mechanisms [36,37], we asked whether a well-known NER substrate would exhibit the same activation of DNA unwinding described above. For this we examined DNA substrates with a single thymine dimer positioned in the middle of a 40 bp duplex (Methods and Supplemental Figure 3). Indeed, in the presence of the thymine dimer in the displaced strand, we observed a 1.6 ± 0.2-fold activation of unwinding (Figure 2B). However, as the prepared substrate only has 86% thymine dimer as judged by ELISA with a thymine-dimer-specific antibody (Methods and Supplemental Figure 3), this represents an actual activation of 1.9 ± 0.2-fold (Figure 2C).

Importantly, we considered whether the effect of the thymine dimer could be due to a reduction in the thermodynamic stability of the duplex. However, this possibility was excluded as the abasic substrate does not increase unwinding (Figure 2A) and both substrates have similarly small reductions in their melting temperatures compared to the fully complementary duplex (Supplemental Figure 4).

### UvrD1 remains in the vicinity of the modification once it is encountered

Since the proximity of a protein often enhances the fluorescence intensity of Cy3 through photoisomerization-related fluorescence enhancement (PIFE) [38,39], we were able to follow the approach of the helicase to the Cy3 modification and subsequent events during unwinding. In the absence of protein, Cy3 fluorescence is constant as a function of time as expected (Supplemental Figure 5). In the presence of UvrD1 and ATP, it exhibits several time-dependent phases (Figure 3).

**Figure 3:**
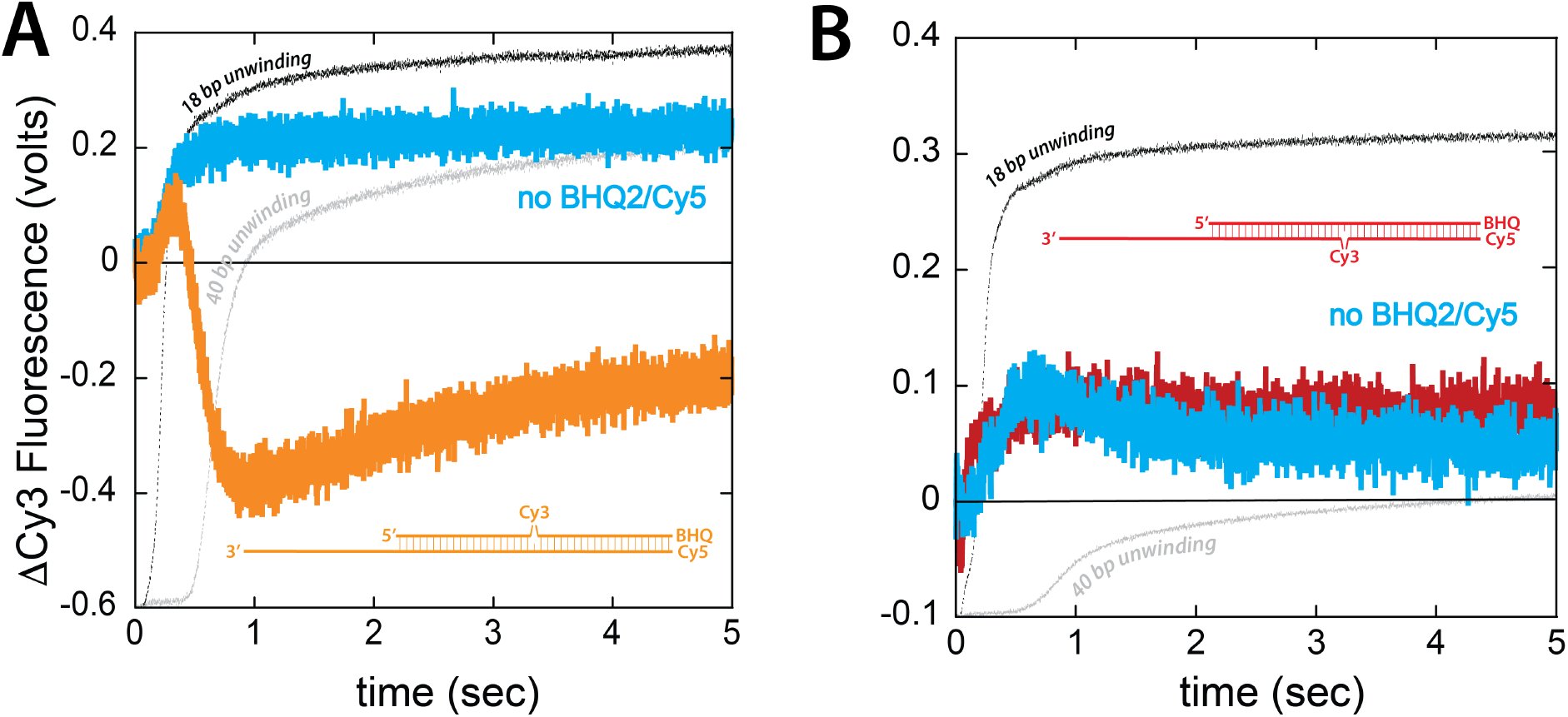
Protein-induced fluorescent enhancements as a function of time on three different 40 bp substrates. **(A)** Changes in Cy3 fluorescence over time in the presence (orange) and absence (cyan) of the downstream BHQ2 for the substrate labeled internally on the displaced strand. **(B)** Changes in fluorescence over time in the presence (red) and absence (cyan) of the downstream BHQ2 for the substrate labeled internally on the loading strand. In both (A) and (B), the unwinding kinetics (taken from figure 1B) are indicated by the thin black (18 bp) and gray (40 bp) lines in the background.

Initially, Cy3 fluorescence increases (Figure 3A, orange). This part of the trace follows the kinetics of the unwinding signal from 18 bp duplex substrates (Figure 3A, black), consistent with the interpretation that the Cy3 fluorescence increase reports on the arrival of enzymes at the position of the modification (20 bp). After reaching a maximum, the fluorescence then decreases with kinetics that coincide with those of the unwinding signal observed for the 40 bp substrate (Figure 3A, grey). However, instead of returning to baseline, the signal decreases significantly below that of the control Cy3 fluorescence (*i*.*e*., negative ΔF). Finally, the signal reaches a minimum and slowly begins to increase again.

We reasoned that the dramatic decrease in Cy3 fluorescence may be caused by the black hole quencher and Cy5 at the 3’ and 5’ downstream ends respectively being brought closer to the Cy3 as the reaction progresses. If this were true, it would suggest that the enzyme stays in the proximity of the Cy3 while it continues to unwind and reel in the downstream DNA. To test this directly, we monitored Cy3 PIFE using substrates that lacked the BHQ2/Cy5 pair used to monitor unwinding (Supplemental Table 1, Cy3-1-NQs). Using this substrate, the Cy3 fluorescence signal initially increases with the same kinetics as before but, after reaching a maximum, sustains that level for the remainder of the reaction (Figure 3A, cyan). Thus, the downstream 3’ end of the displaced strand must be approaching the Cy3 and UvrD1 must maintain contact with the Cy3 as it continues to unwind the downstream duplex.

Lastly, we looked at the PIFE signal on the substrate with Cy3 positioned in the middle of the 40 bp duplex in the loading strand, which reduces unwinding (Figure 1A, red). Here, the fluorescence again increases over time consistent with the time required for the helicase to arrive at the Cy3 position (Figure 3B, red). However, upon reaching a maximum, the signal remains at a constant level suggesting that the helicase is stuck or paused at the modification and that, in these complexes, the downstream 3’ end of the unwound strand is not brought closer to the Cy3. Consistently, the removal of the BHQ and Cy5 from the downstream end does not significantly change the kinetics of the observed fluorescent enhancement (Figure 3B, cyan).

### A decrease in helicase dissociation rate leads to a transient increase in helicase processivity

Processivity (P) is a property of processive motors that describes the average number of forward steps taken per binding event prior to dissociation and can be quantified as the ratio of the translocation rate to the sum of the translocation and dissociation rates. We postulated that a possible explanation for the increased amount of unwinding of the 40 bp substrate was that the modifications on the displaced strand result in increased processivity. More specifically, as the lag phase is not affected by the presence of the modification, this increase in processivity would have to be due to a decrease in the dissociation rate as opposed to an increase in the translocation rate (Figures 1 and 2). Experimentally, processivity can be estimated from the change in the fraction of DNA unwound as a function of dsDNA length. Thus, to test this idea, we measured the fraction of DNA unwound on substrates with increasing duplex length in the presence and absence of the PEG modification in the displaced strand (Figure 4 and Supplemental Figure 6).

**Figure 4:**
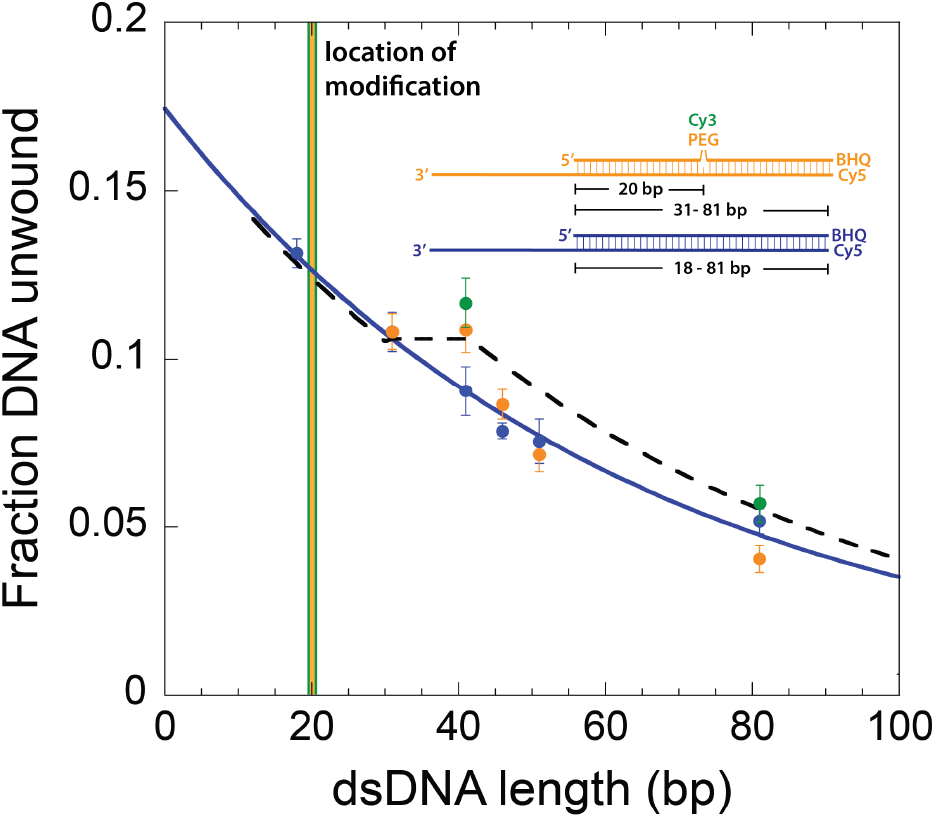
Activation via temporarily increased processivity. The fraction unwound on unmodified (blue) and substrates modified with PEG (orange) or Cy3 (green) at position 20 as a function of DNA length. The increased fraction unwound due to the modification is limited to a range of duplex lengths between 30-40 bp following the modification consistent with a temporary increase in processivity.

The fraction of DNA molecules unwound as a function of the duplex length of a series of unmodified substrates of 18, 31, 41, 46, 51, and 81 bp were used to establish a baseline processivity (Methods, Figure 4, blue, P = 0.98). Modified substrates contained a PEG moiety 20 bp downstream (in the direction of unwinding) from the DNA junction on the displaced strand followed by 11, 21, 26, 31, or 61 bp (Figure 4, orange). The duplex length dependence exhibited several interesting features. First, the fraction of modified DNA substrates unwound was the same as for the unmodified DNA for the 31 bp duplex even though the modification is located upstream at the 20^th^ base-pair. Second, the 41 bp modified substrate exhibited the same fraction of DNA molecules unwound as the 31 bp substrate suggesting that all the enzymes that reach 31 bp also make it to 41 bp. Third, the difference between the modified and unmodified DNA diminishes for longer substrates. Data acquired with 41 bp and 81 bp with a Cy3 modification in the same positions as the PEG exhibited a similar trend (Figure 4, green).

Based on the 10 bp shift of the onset of the effect relative to the position of the modification, we conclude that the interaction between the modification and the helicase that results in reduced dissociation takes place in the unwound, single-stranded DNA state. To understand the complex behavior that follows, we compared the data to theoretical expectations derived from different models. Since the effect of the modification disappears for longer lengths, any effect on the dissociation rate must be transient. The data do not follow the simplest model, in which the dissociation rate returns to its value on unmodified DNA after reaching 41 bp (Figure 4, dashed line). We next tried models where, after 41 bp, the dissociation rate increases beyond its basal value. While these classes of model were better able to account for different phases of the data (Supplemental Figure 7), a single dissociation rate does not fit the shape of the entire length dependence suggesting that the enzyme transitions through multiple states after encountering the modification, each with different properties.

Taken together, our analyses suggest that when UvrD1 encounters a modification in the displaced strand, there is an abrupt, but short-lived, decrease in dissociation rate. The overall result is that DNA unwinding is enhanced over a specific range of lengths 10 – 20 bp downstream from the modification.

### Displaced strand thymine dimer activates *E. coli* UvrD dimers

Given that our recent work demonstrates that UvrD1 family members dimerize using the same 2B-2B subdomain interface [20,21,40], we asked whether the effects of modifications placed in the 5’-3’ displaced strand would similarly affect the unwinding by *Ec*UvrD dimers and performed measurements on the 40 bp thymine dimer substrates (Methods). Indeed, using 90 nM protein and 10 nM DNA, we also observe enhancement of *Ec*UvrD unwinding activity (Figure 5, Methods). After correction for the fraction of modified DNA, this is equivalent to a 1.9 ± 0.3-fold increase which is of the same order of magnitude observed with *Mtb* UvrD1. Therefore, we posit that the activation of dimer unwinding by displaced-strand modifications is a property shared across the UvrD family of helicases.

**Figure 5:**
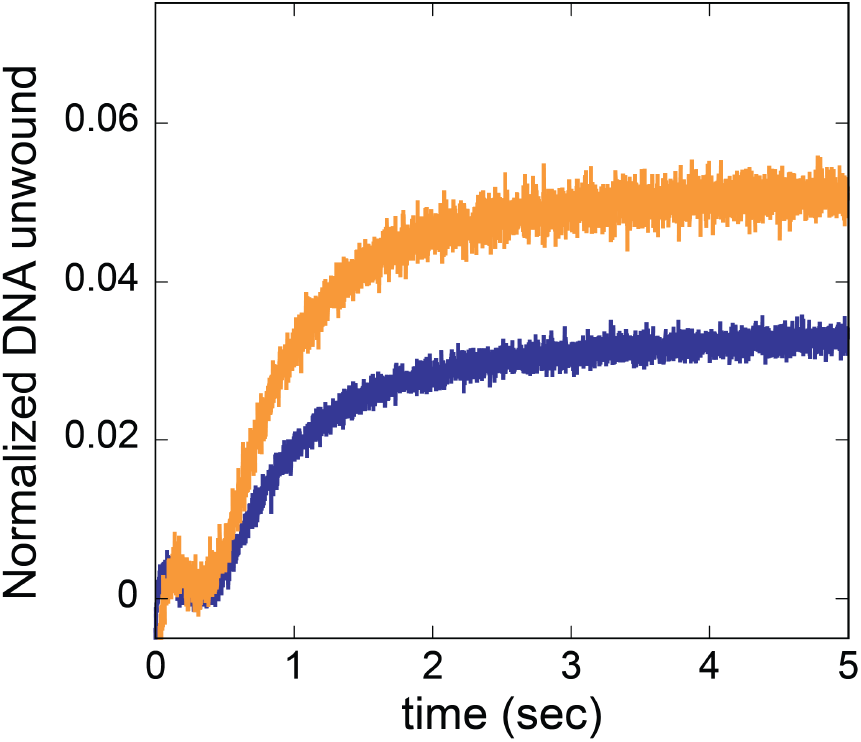
Activation of DNA unwinding by E. coli UvrD. A thymidine dimer in the displaced strand (orange) enhances unwinding compared to a 40 bp unmodified DNA (blue). Data are normalized to fraction unwound of an 18 bp unmodified DNA by E. coli UvrD (Supplemental Figure 8).

## Discussion

DNA unwinding by UvrD-family helicases requires dimerization in the absence of other activating factors, but the mechanism dimer unwinding is not yet fully clear. Structures of dimeric UvrD1 bound to a DNA junction show a dramatic repositioning of the 2B subdomain away from its contacts with duplex DNA observed in monomeric structures [20]. This indicates that the 2B:dsDNA interactions are inhibitory and that at least part of the mechanism of activation involves removal of this inhibition. In fact, the structure shows limited dsDNA contacts with two subunits bound one in front of the other to the single-stranded DNA loading strand. The open question that remains is how the individual subunits move relative to the DNA and relative to each other during strand separation.

Models may include the independent or coupled translocation of each subunit along the loading strand with or without exchanging register. For example, a force-dependent study of dimer catalyzed DNA unwinding suggested a scrunching inch-worm model where each subunit translocates along the single-stranded DNA of the loading strand which is compacted between the two subunits during each step of the helicase [33]. However, another class of models not excluded by previous observations include interactions with the displaced strand. We set out to exclude this specific class of models by asking whether modifications in the displaced strand affect unwinding. In contrast, our data suggest that the opposite is true and that UvrD1 makes significant, and potentially mechanistically important, interactions with the displaced strand.

While the structure of dimeric UvrD1 bound to a DNA junction shows the two subunits bound to the loading strand [20], caution must be taken in interpreting this conformation. The structure represents an initially bound complex prior to unwinding as no ATP was present. Given the evidence of interactions with the displaced strand presented here, the question pivots to the nature of those interactions during active unwinding. Our length-dependent data (Figure 4) reveal that the enhancement occurs only at lengths past 30 bp in that the 30 bp modified and unmodified lengths were unwound equally and only at duplex lengths of 40 bp and longer do we see the effect. Considering that the modifications were located at 20 bp, we infer that the causal interaction occurs after unwinding and in the context of a single-stranded displaced strand. A concrete structural explanation for the interaction remains to be discovered.

A further aspect of mechanism to be considered is whether individual subunits contribute different functionality during the entirety of a processive DNA unwinding reaction. This question is often posed as whether the complex proceeds along the DNA via an inchworm or a rolling mechanism [33,41,42]. Each could be imagined regardless of whether the enzyme interacts with one or both strands. One could even imagine a mechanism where these two limiting pictures co-exist in a distributed manner (*i*.*e*., some processive inchworming interspersed with the switching of lead/trailing subunits). However, biochemical investigations of the outcomes of mixing mutant *E. coli* UvrD suggest that perhaps each subunit takes on a specific role during the reaction and that these roles can be combined to generate activity. Specifically, the W256A mutant UvrD is incapable of unwinding even in its dimeric form but supports unwinding in the context of hybrid dimers with WT subunits, suggesting a division of roles that the tryptophan mutant effectively separates [30]. This indicates a constant role for each subunit (dependent perhaps on the initial binding position) and, in the context of our model, suggests that one subunit interacts with the loading strand and one with the displaced strand during a processive inchworm type of unwinding reaction. This would, in turn, imply that the structure of an actively unwinding UvrD complex will be distinct from the observed initially bound structure.

Biologically, the implications of our results center around the role of UvrD1 and related enzymes in DNA repair. Specifically, during bacterial nucleotide excision repair UvrDs are responsible for catalyzing the removal of the damaged strand after identification by UvrAB and the introduction of nicks surrounding the lesion by UvrC [43,44]. Our results are particularly suggestive as lesions being removed by this process map directly to what would be the displaced strand in our experiments. Furthermore, the thymine dimer, a biologically relevant lesion, shows the most dramatic stimulation even though it represents a smaller structural perturbation. This raises the intriguing possibility that the presence of lesions stimulates UvrD activity and leads to more efficient and specific NER. One caveat to this idea is that UvrDs are also activated by auxiliary factors [12,45–47] and it is not known which *in vivo* processes depend on dimerization [48]. An unexplored further possibility is that dimerization, and the interaction with damaged displaced strands we describe here, may play roles in non-NER pathways [49].

## Materials and Methods

### Protein and nucleic acid preparation

Wild-type (WT) Mtb UvrD1 and C451A mutant were overexpressed and purified as previously described [19] with the following modification: Size exclusion chromatography was not used for the WT protein as it was adequately pure after elution from the heparin column. Both proteins were dialyzed against storage buffer (50 mM Tris-HCl, pH 8.0, 200 mM NaCl, and 25% (v/v) glycerol), spin concentrated to a concentration between 12-25 μM, aliquoted and flash frozen with liquid nitrogen for storage at −80 °C. The protein concentration was determined by absorbance spectroscopy at 280 nm using a molar extinction coefficient of 63385 M^−1^cm^−1^ determined from the sequence. WT *E. coli* (*Ec*) UvrD protein was overexpressed and purified as previously described [50]. *Ec*UvrD concentration was determined by absorbance spectroscopy at 280 nm using a molar extinction coefficient of 1.06 x 10^5^ M^−1^cm^−1^.

The single-stranded DNA oligos used to assemble the various double-stranded DNA substrates were synthesized by Sigma-Millipore and oligos containing blackhole quencher 2 (BHQ2) were synthesized by IDTdna. Supplemental Tables (1-3) list each DNA substrate, the oligo sequences and the calculated extinction coefficient used to determine the annealed, nucleic acid concentration. The DNA substrates were designed with 3’ 20 nt deoxythymidilate tails to allow for helicase loading [16,19]. The 5’-end of the loading strand is labeled with Cy5 while the 3’-end of the complementary strand is labeled with BHQ2. While the duplex is annealed, BHQ2 quenches the Cy5 fluorescence. Unwinding of the DNA duplex results is release of the BHQ2 labeled strand, resulting in an increase in the Cy5 fluorescence [19,46]. The DNA substrates were prepared by annealing the loading strand with its complement strand (1:1.02, slight molar excess) in 1x annealing buffer (10 mM Tris-HCl, pH 8.0, 50 mM NaCl). The DNA solution was heated at 95 °C for 5 minutes, then slowly cooled to 25 °C over a 4-hour period. The annealed DNA was verified by native polyacrylamide gel electrophoresis (10% gel in 1x TBE buffer, ran at 120 V) and stored at −20 °C. Preparation of the thymine dimer containing DNA substrate and substrates longer than 50 bp are discussed in supplemental methods. Molar extinction coefficients at 260 nm were calculated as in [51] using equations 1 and 2 and molar extinction coefficients (M^−1^ cm^−1^) for different components as follows: 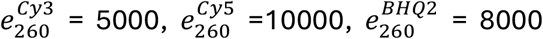, and 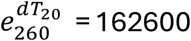.

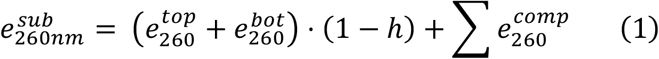

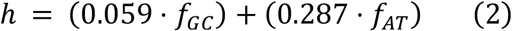

Where 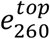 and 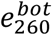 are the extinction coefficients at 260 nm for the complementary top and bottom ssDNA oligos, respectively, that form the duplex region; *h is the hypochromicity;* 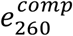 is the extinction coefficient at 260 nm for present components and *f*_*GC*_ and *f*_*AT*_ are the fraction GC and AT, respectively, for the duplex region.

### Stopped-flow kinetic measurements

All stopped-flow kinetic measurements were conducted at 25 °C using an Applied Photophysics SX20-LED stopped-flow instrument with 1:1 mixing in a 100 ul total shot volume and 1 ms deadtime. All measurements with UvrD1 were carried out in buffer containing: 20 mM TRIS-HCl, pH 8.0 (25 °C), 75 mM NaCl, and 20% (v/v) glycerol. Experiments with *Ec*UvrD were carried out in buffer containing: 10 mM TRIS-HCl, pH 8.3 (25 °C), 20 mM NaCl, and 20% (v/v) glycerol. All reported concentrations are the final concentrations after mixing. Samples were loaded into the stopped-flow instrument and incubated for 5 mins at 25 °C before initiation. After initiation, the reaction was monitored for 10 sec. A reaction vol of 1 mL was sufficient to obtain 10 traces of which 6-8 were averaged together.

#### Fluorescence DNA unwinding assay

The single-round kinetics of DNA unwinding were monitored by following the increase in Cy5 fluorescence upon release of the quencher-labeled complementary strand. A 625 nm LED was used to excite Cy5 and its fluorescence emission was detected at wavelengths >650 nm with a long pass filter. Reactions were initiated by rapidly mixing a solution of helicase pre-incubated with DNA substrate with a solution containing MgCl_2_, ATP, and excess ssDNA trap. The ssDNA trap functions in two ways: (1) It is complementary to the unwound strand, preventing the unwound DNA from reannealing and (2) It binds to free UvrD1, preventing UvrD1 from reinitiating DNA unwinding, thus establishing the single-round conditions. Reactions containing UvrD1 have 5 mM MgCl_2_, 1 mM ATP, and 10 μM ssDNA trap. Reactions containing *Ec*UvrD have 2 mM MgCl_2_, 1 mM ATP, 1 μM ssDNA trap and 4 μM hairpin trap, that binds *Ec*UvrD with higher affinity [16]. Positive and negative control traces were collected to convert the fluorescence signal into fraction DNA unwound as previously described [19]. Unwinding traces where then normalized to the maximum DNA unwound for the 18 bp control substrate to account for any variation in prep-to-prep dimer fraction. The fold enhancement in DNA unwound was calculated by taking the ratio of fraction DNA unwound on modified and unmodified substrates.

DNA unwinding processivity of the active helicase fraction was determined by fitting the fraction of DNA unwound as a function of unmodified dsDNA length using equation (3) modified from [52] to account for finite active helicase fraction.

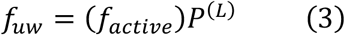

Where *f*_*active*_ is the active helicase fraction, *P* is the unwinding processivity, and is *L* is the dsDNA length (bp).

#### Helicase arrival PIFE assay

DNA substates modified with Cy3 can detect the presence of UvrD1 through protein induced fluorescence enhancement (PIFE) allowing us to monitor the arrival of UvrD1 at the Cy3 modification during single-round DNA unwinding. A 535 nm LED with a 550 nm short-pass cutoff filter was used to excite Cy3 and its fluorescence emission was detected at wavelengths >570 nm with a long pass filter. Reactions were initiated as in the DNA unwinding assays described above. A negative control trace was collected by rapidly mixing a DNA only solution with a solution containing MgCl_2_, ATP, and excess ssDNA trap. The negative control was subtracted from the trace containing UvrD1, yielding the net Cy3 fluorescence change during DNA unwinding.

### Simulation of DNA unwinding time courses

To model the fraction DNA unwound as a function of dsDNA length in the presence and absence of PEG modification we simulated single-round DNA unwinding time courses in MatLab (Mathworks) using the following scheme:

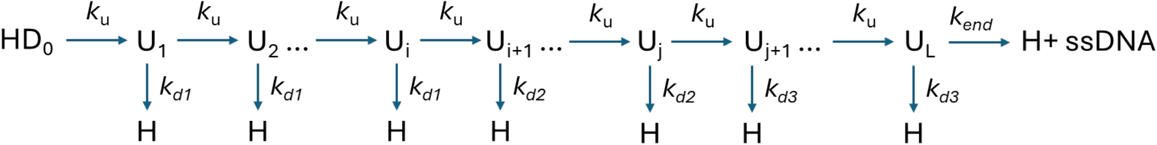

where an active helicase fraction initiates DNA unwinding taking i steps at a rate k_u_ (60 bp/sec). Upon encountering the modification, the dissociation rate changes from k_d1_ (1 s^−1^) to k_d2_ (k_d2_ = 0 s^−1^). After j additional steps, the dissociation rate either returns to the intrinsic value (k_d3_ = k_d1_) or becomes faster (k_d3_ > k_d1_)for the remainder of the unwinding steps up to the length of the DNA substrate (L). Helicase dissociation and release of ssDNA occur with a rate k_end_.

We simulated DNA unwinding time courses for L = 10-100 bp varying the position where the modification takes effect (5, 8,10 bp past actual modification position (20 bp from ss/dsDNA junction)) and for different values of k_d3_ (k_d3_ = 1 s^−1^ – 5 s^−1^) to try to reproduce the observed PEG-dependent length dependence of the fraction of DNA unwound.

## ACKNOWLEDGEMENTS

We thank Dr. Tim Lohman for many fruitful discussions that helped guide this work from beginning to end and Dr. Binh Nguyen (Lohman lab) for providing WT *E. coli* UvrD.

## AUTHOR CONTRIBUTIONS

E.J.T. and E.A.G. designed research; E.J.T. performed experiments and analyzed the data. A.C. provided initial preparations of UvrD1. E.J.T. and E.A.G. wrote the paper. E.A.G. guided the project.

