## Supplemental Materials and Methods for "Bulky lesions on the displaced strand stimulate DNA unwinding by dimeric UvrD-family helicases"

**Construction of modified DNA unwinding substrates longer than 51bp**

Due to limitations in commercial synthesis of modified DNA oligos with the black hole quencher (BHQ2) longer than 50 nts, we constructed longer unwinding substrates by annealing two separate duplexes with complementary overhangs then ligated them together with T4 DNA ligase as shown in the diagram below.

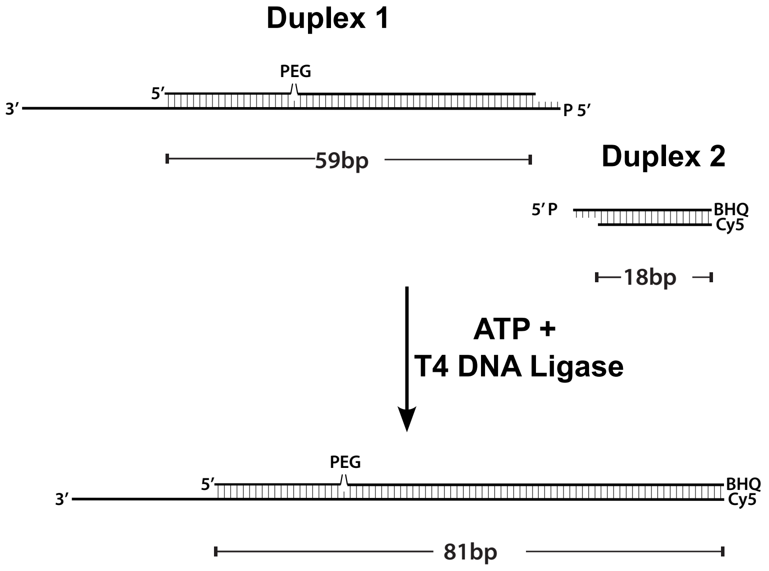


Duplex 1 has a 3' tail region for loading helicase with or without the PEG modification and a 5'-phosphorylated, 4-base overhang for ligation to duplex 2. Duplex 2 is a short duplex with the BHQ2 and Cy5 labels that has a 5'-phosphorylated, 4-base overhang. In a 50 µL volume of 1x T4 DNA ligase buffer, 5 µM of each duplex was combined with 1600 U of T4 DNA ligase (NEB). The reaction was incubated at 25 °C for 2 hrs, then at 16 °C overnight. Supplemental Table 4 lists all oligos used for duplexes 1 and 2 to construct the 81 bp PEG, Cy3, and control substrates highlighted in Supplemental Table 1 and 2.

Full-length ligation products were gel purified by native PAGE (10%, 1x TBE, 120V), were the slowest migrating species identified by UV shadowing, and clearly exhibited a color consistent with the presence of Cy5/BHQ2 labels (purplish). The DNA was electro-eluted (Elutrap system) from the excised gel slices into a 250-300 µl volume (1x TBE, 120V) and was subsequently concentrated by ethanol precipitation as follows. Prior to precipitation the NaCl was adjusted to 0.3 M and 2.5x vol of ice-cold Ethanol was added to precipitate the DNA. To facilitate precipitation, the solutions were placed at -80 °C for 30 min, then centrifuged at 13000 rpm for 30 min at 4 °C to pellet the DNA. The DNA pellet was washed with 70% ethanol, pelleted by centrifugation, air dried, and resuspended in 1x anneal buffer (10 mM TRIS-HCl, pH 8.3 (25 °C), 50 mM NaCl). Final products were confirmed by native PAGE (10%, 1x TBE, 120V).

**Preparation of UV induced thymine dimer modification**

Since commercially available synthesis of oligos containing the cis-syn cyclobutane thymine dimer modification was cost prohibitive for our study we devised a strategy to generate a DNA substrate containing the thymine dimer modification induced by UV irradiation. We note this method will generate a mixture of thymine dimers, both the cis-syn cyclobutane thymine dimer and the (6-4) thymine dimer photoproduct. We did not attempt to separate these products but used an immuno-absorbent assay and absorption spectroscopy to quantify their amounts. Our measurements indicate ~86% of the DNA has a thymine dimer modification.

To minimize UV exposure of the full DNA substrate, only the non-loading strand was irradiated with UV. The oligo was designed to have a single region in the middle with three adjacent pyrimidines (Supplemental Table 3). A thymine dimer can form between the first two or last two deoxythymines. A 25 µM solution of the oligo was prepared in 1x anneal buffer. To irradiate the sample, 12 µL of oligo solution was loaded into a quartz cuvette with a 0.010 cm pathlength and placed at 5 °C on a reflective tray. Lower temperature facilitates dimer formation by reducing thermal motion of the bases [1]. A UV (254 nm) lamp (Model UVGL-25: 15 vols, 60 Hz, 0.16 AMPS) was used to irradiate the sample and was placed 6 mm from the surface of the cuvette. At this distance the UV source produces 1.92 $kJ/m^2/min$ as measured by a UVX digital radiometer. The sample was irradiated for 30 minutes. Longer irradiation times resulted in significant loss of the BHQ2 absorption band between 450-700 nm (Supplemental Figure 2B). Oligos exposure for 30 min, with and without BHQ2, showed a decrease in absorption at 260 nm, as expected for syn cyclobutane thymine dimer formation, and an increase in absorption at 325 nm, as expected for dimer conversion to the (6-4) dimer photoproduct (Supplemental Figure 2A) [1]. The concentration of the (6-4) photoproduct was determined at 325 nm using a molar extinction coefficient of 4600 $M^{-1}cm^{-1}$ [2] and indicated 22% of the oligo has the (6-4) photoproduct. The presence of the cis-syn cyclobutane thymine dimer was confirmed by EMSA using a cis-syn cyclobutane thymine dimer specific antibody [3]. Supplemental Figure 2C shows a portion of the UV treated oligo band is shifted in the presence of the antibody while over exposure leads to loss in the shifted band, most likely due to conversion to the (6-4) photoproduct which is not recognized by the antibody [3]. The DNA unwinding substrate was prepared by annealing the UV-treated oligo with the loading strand oligo listed in Supplemental Table 3 as described in the main text.

*Cis-syn cyclobutane thymine dimer immuno-absorbent assay*. An immuno-absorbent assay was performed to quantify the cis-syn cyclobutane thymine dimer present in the oligo. The UV treated oligo was annealed to a 5'-biotin labeled 15nt oligo to allow for DNA binding to a streptavidin coated ELISA plate (Pierce, cat#: 15407) as depicted in the diagram below.

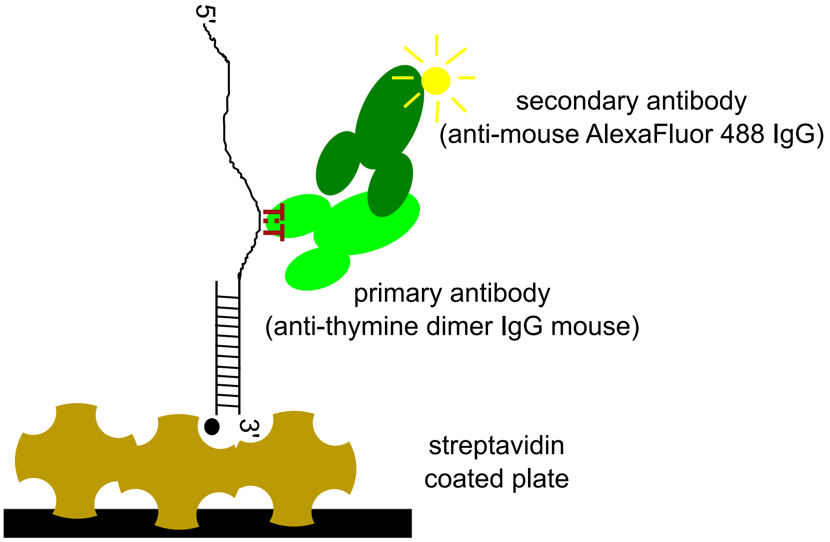


The region of the DNA containing the thymine dimer is single-stranded to allow for anti-thymine dimer monoclonal antibody recognition (Millipore-Sigma, cat#: T1192). The presence of the anti-thymine dimer antibody (mouse) was detected by a secondary anti-mouse antibody labeled with Alexa Fluor 488 (Thermo-Fischer (Invitrogen), cat#: A10680).

The ELISA plate manufacturer's protocol was followed to absorb the DNA and antibodies for detection as follows. Wells of the streptavidin-coated plate were washed three times with 100 µL wash buffer (25 mM TRIS-HCl pH 7.5 (25 C), 150 mM NaCl, 0.1% acetylated BSA (Millipore-Sigma, cat#: B2518), and 0.05% Tween-20) with the buffer being removed by aspiration. A 100 nM solution (50 µL) of thymine dimer, biotinylated DNA duplex in wash buffer was added to wells in triplicate for each primary antibody dilution and allowed to incubate at 25 °C for 2 hrs. Post incubation, solution was collected by pipette and the remaining concentration in solution measured by absorbance, indicating 80 +/- 2 nM of DNA remained bound in the wells. Wells were washed three times with 100 µL wash buffer, then a 1:50 (267 nM) and 1:25 (533 nM) dilution of primary antibody in wash buffer (50 µL) was added to wells containing the thymine dimer DNA and to wells without DNA and incubated for 30 min at 25 °C. The primary antibody solution was removed, and the wells washed three times with 100 µL wash buffer. A 300 nM solution of AlexaFluor 488 secondary antibody in wash buffer (50 µL) was added to each well and incubated for 30 min at 25 °C. Post incubation, the wells were washed three times with wash buffer (100 µL), refilled with 100 µL wash buffer and scanned in a CLARIOstar$^{Plus}$ fluorescent plate reader (BMG LabTech) at 25 °C. The plate was scanned in fluorescence intensity mode with monochromator adjusted 488 nm excitation (14 nm bandwidth) and 535 nm emission (30 nm bandwidth) using a 507 nm dichroic top mirror to filter out excitation light.

A standard curve of Alexa Fluor 488 surface bound antibody was constructed using a biotin labeled mouse secondary antibody (Millipore-Sigma (Invitrogen), cat#: 31824,). Wells were washed and treated as above with ~240 nM solution of primary antibody. A serial dilution of AlexaFluor 488 secondary antibody was added to the wells, incubated for 30 min at 25 °C, washed, and prepared for scanning in the fluorescent plate reader as described above. After accounting for non-specific binding to the well surface, the fluorescence was linear over 80 nM range of secondary antibody (Supplemental Figure 2). Under the buffer conditions, fluorescence due to non-specific binding was 4% or less and was subtracted before determining the amount of thymine dimer present. Based on the standard curve, the observed fluorescence in the presence of anti-thymine dimer antibody indicates 64% of the bound oligo has the cis-syn cyclobutane thymine dimer.

**DNA melting curve analysis of thymine dimer substrate**

We measured the thermal stability of the thymine dimer substrate and abasic site substrates with a thermal melt monitoring the absorbance of the nucleic acid at 260 nm as the temperature was changed. DNA substrates were prepared as described in methods with oligos lacking the Cy5 and blackhole quencher 2 dyes. The modified substrates along with control substrates lacking the modification were diluted to an absorbance of 0.3 in buffer containing 20 mM $NaPO_4$, 75 mM NaCl, and 20% (v/v) glycerol. The Tris buffer was changed to phosphate to minimize changes in pH as the temperature was changed. The thermal melt was carried out in a Cary Agilent UV-vis Peltier 3500 spectrophotometer with 2 °C change/ min from 15C to 95 °C with absorbance measurements was taken at each degree C. The melting curves were buffer (blank) corrected and the first derivative was calculated to determine the melting temperature ($T_m$).

**Supplemental Table 1: Unmodified and Cy3-modified substrates.**

Highlighted sequences were constructed from individual oligos listed in Supplemental Table 4.

| Name | Substrate Design | Extinction Coefficient ($M^{-1}cm^{-1}$) | Oligo Sequences |
|---|---|---|---|
| Con18 | 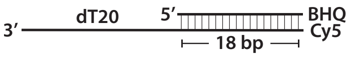  | 466785.36 | 5'GCCCTGCTGCCGACCAAC***BHQ2***-3'<br><br>5'***CY5***GTTGGTCGGCAGCAGGGCTTTTTTTTTTTTTTTTTTTT3' |
| Con41 | 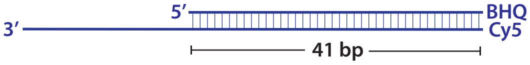  | 824107.10 | 5'GCCCTGCTGCCGACCAACGATTGGTTACATTCCCGCTGC TG***BHQ2***-3'<br><br>5'***CY5***CAGCAGCGGGAATGTAACCAATCGTTGGTCGGCAGC AGGGCTTTTTTTTTTTTTTTTTTTT3' |
| Cy3-1 | 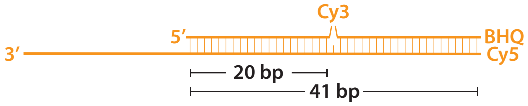  | 830632.55 | 5'GCCCTGCTGCCGACCAACGA***CY3***TGGTTACATTCCCGCT GCTG***BHQ2***-3'<br><br>5'***CY5***CAGCAGCGGGAATGTAACCAATCGTTGGTCGGCAGC AGGGCTTTTTTTTTTTTTTTTTTTT3' |
| Cy3-2 | 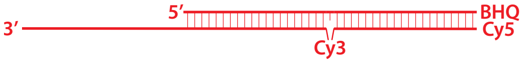  | 824670.4 | 5'GCCCTGCTGCCGACCAACGATTGGTTACATTCCCGCTGC TG***BHQ2***-3'<br><br>5'***CY5***CAGCAGCGGGAATGTAACCA***CY3***TCGTTGGTCGGCA GCAGGGCTTTTTTTTTTTTTTTTTTTT3' |
| Cy3-3 | 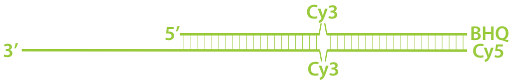  | 822669.2 | 5'GCCCTGCTGCCGACCAACGA***CY3***TGGTTACATTCCCGCT GCTG***BHQ2***-3'<br><br>5'***CY5***CAGCAGCGGGAATGTAACCA***CY3***TCGTTGGTCGGCA GCAGGGCTTTTTTTTTTTTTTTTTTTT3' |
| Cy3-5 | 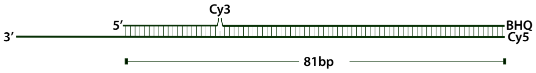  | 1468143.82 | 5'GCCCTGCTGCCGACCAACGA***Cy3***TGGTTACATTCCCGCT GCTGGAGAAACGGCCGAATACATGGAGAAACGGCCGAATA CACC***BHQ2***-3'<br><br>5'***CY5***GGTGTATTCGGCCGTTTCTCCATGTATTCGGCCGTTTC TCCAGCAGCGGGAATGTAACCAATCGTTGGTCGGCAGCAG GGCTTTTTTTTTTTTTTTTTTTT |
| Cy3-1-NQs | 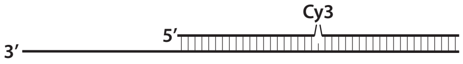  | 812632.55 | 5'GCCCTGCTGCCGACCAACGA***CY3***TGGTTACATTCCCGCT GCTG3'<br><br>5'CAGCAGCGGGAATGTAACCAATCGTTGGTCGGCAGCAG GGCTTTTTTTTTTTTTTTTTTTT3' |
| Cy3-2-NQs | 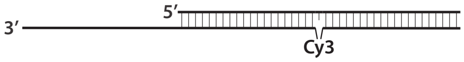  | 767923 | 5'GCCCTGCTGCCGACCAACGATTGGTTACATTCCCGCTGC TG3'<br><br>5'CAGCAGCGGGAATGTAACCA***CY3***TCGTTGGTCGGCAGCA GGGCTTTTTTTTTTTTTTTTTTTT3' |
| Cy3-3-NQs | 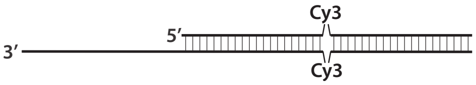  | 766360.4 | 5'GCCCTGCTGCCGACCAACGA***CY3***TGGTTACATTCCCGCT GCTG3'<br><br>5'CAGCAGCGGGAATGTAACCA***CY3***TCGTTGGTCGGCAGCA GGGCTTTTTTTTTTTTTTTTTTTT3' |

**Supplemental Table 2: Polyethylene glycol-modified substrates.**

Highlighted sequences were constructed from Individual oligos listed in Supplemental Table 4.

| Name | Substrate Design | Extinction Coefficient ($M^{-1}cm^{-1}$) | Oligo Sequences |
|---|---|---|---|
| PEG-1 | 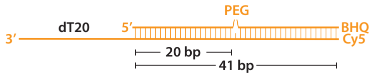 | 825732.55 | 5'GCCCTGCTGCCGACCAACGA***Sp9***TGGTTACATTCCCGCTGCTG***BHQ2***-3'<br><br>5'***CY5***CAGCAGCGGGAATGTAACCAATCGTTGGTCGGCAGCAGGGCTTTTTTTTTTTTTTTTTTTT3' |
| PEG-2 | 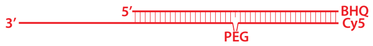 | 826588.05 | 5'GCCCTGCTGCCGACCAACGATTGGTTACATTCCCGCTGCTG***BHQ2***-3'<br><br>5'***CY5***CAGCAGCGGGAATGTAACCA***Sp9***TCGTTGGTCGGCAGCAGGGCTTTTTTTTTTTTTTTTTTTT3' |
| PEG-3 | 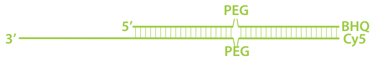 | 819658.5 | 5'GCCCTGCTGCCGACCAACGA***Sp9***TGGTTACATTCCCGCTGCTG***BHQ2***-3'<br><br>5'***CY5***CAGCAGCGGGAATGTAACCA***Sp9***TCGTTGGTCGGCAGCAGGGCTTTTTTTTTTTTTTTTTTTT3' |
| PEG-4 | 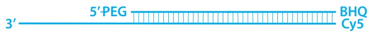 | 830822.25 | 5'***Sp9***GCCCTGCTGCCGACCAACGATTGGTTACATTCCCGCTGCTG***BHQ2***-3'<br><br>5'***CY5***CAGCAGCGGGAATGTAACCAATCGTTGGTCGGCAGCAGGGCTTTTTTTTTTTTTTTTTTTT3' |
| PEG-5 | 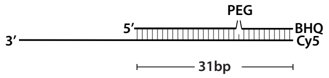 | 656300.81 | 5'GCCCTGCTGCCGACCAACGA***Sp9***TGGTTACATT**BHQ2**-3'<br><br>5'***CY5***AATGTAACCAATCGTTGGTCGGCAGCAGGGCTTTTTTTTTTTTTTTTTTTT3' |
| PEG-6 | 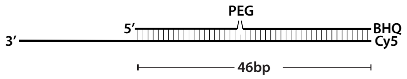 | 905139.48 | 5'GCCCTGCTGCCGACCAACGA***Sp9***TGGTTACATTCCCGCTGCTGGAGAA***BHQ2***-3'<br><br>5'***CY5***TTCTCCAGCAGCGGGAATGTAACCAATCGTTGGTCGGCAGCAGGGCTTTTTTTTTTTTTTTTTTTT3' |
| PEG-7 | 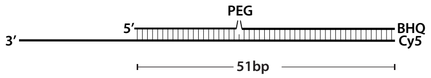 | 984125.58 | 5'GCCCTGCTGCCGACCAACGA***Sp9***TGGTTACATTCCCGCTGCTGGAGAAACGGC***BHQ2***-3'<br><br>5'***CY5***GCCGTTTCTCCAGCAGCGGGAATGTAACCAATCGTTGGTCGGCAGCAGGGCTTTTTTTTTTTTTTTTTTTT3' |
| PEG-8 | 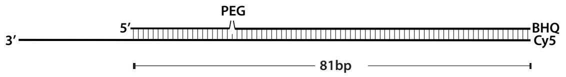 | 1463143.82 | 5'GCCCTGCTGCCGACCAACGA***Sp9***TGGTTACATTCCCGCTGCTGGAGAAACGGCCGAATACATGGAGAAACGGCCGAATACACC***BHQ2***-3'<br><br>5'***CY5***GGTGTATTCGGCCGTTTCTCCATGTATTCGGCCGTTTCTCCAGCAGCGGGAATGTAACCAATCGTTGGTCGGCAGCAGGGCTTTTTTTTTTTTTTTTTTTT |
| Con31 | 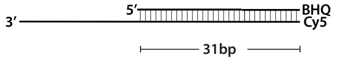 | 663088.16 | 5'GCCCTGCTGCCGACCAACGATTGGTTACATT**BHQ2**-3'<br><br>5'***CY5***AATGTAACCAATCGTTGGTCGGCAGCAGGGCTTTTTTTTTTTTTTTTTTTT3' |
| Con46 | 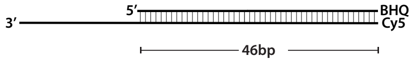 | 909471.75 | 5'GCCCTGCTGCCGACCAACGATTGGTTACATTCCCGCTGCTGGAGAA***BHQ2***-3'<br><br>5'***CY5***TTCTCCAGCAGCGGGAATGTAACCAATCGTTGGTCGGCAGCAGGGCTTTTTTTTTTTTTTTTTTTT3' |
| Con51 | 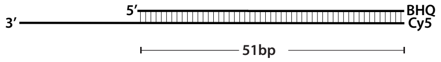 | 981536.5 | 5'GCCCTGCTGCCGACCAACGATTGGTTACATTCCCGCTGCTGGAGAAACGGC***BHQ2***-3'<br><br>5'***CY5***GCCGTTTCTCCAGCAGCGGGAATGTAACCAATCGTTGGTCGGCAGCAGGGCTTTTTTTTTTTTTTTTTTTT3' |
| Con81 | 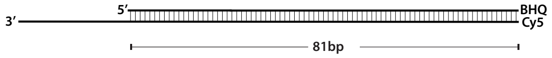 | 1463143.82 | 5'GCCCTGCTGCCGACCAACGATTGGTTACATTCCCGCTGCTGGAGAAACGGCCGAATACATGGAGAAACGGCCGAATACACC***BHQ2***-3'<br><br>5'***CY5***GGTGTATTCGGCCGTTTCTCCATGTATTCGGCCGTTTCTCCAGCAGCGGGAATGTAACCAATCGTTGGTCGGCAGCAGGGCTTTTTTTTTTTTTTTTTTTT |

### Supplemental Table 3: Other modified substrates

| Name | Substrate Design | Molar Extinction Coefficient ($M^{-1}cm^{-1}$) | Oligo Sequences |
|---|---|---|---|
| Ab-1 | 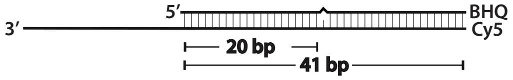 | 814573.2 | 5’GCCCTGCTGCCGACCAACGA***dSp***TGGTTACATTCCCGCTGCTG***BHQ2***-3’<br><br>5’***CY5***CAGCAGCGGGAATGTAACCAATCGTTGGTCGGCAGCAGGGCTTTTTTTTTTTTTTTTTTTT3’ |
| Biotin-1 | 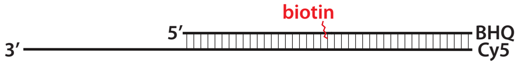 | 824107.1 | 5’GCCCTGCTGCCGACCAACGA***iBiodT***TGGTTACATTCCCGCTGCTG**BHQ2**-3’<br><br>5’***CY5***CAGCAGCGGGAATGTAACCAATCGTTGGTCGGCAGCAGGGCTTTTTTTTTTTTTTTTTTTT3’ |
| ConTT<br><br>TDimer | 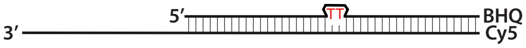 | 829852.62 | 5’CGCGGACGCGAGGACACG**TTT**GGTGCAGGTGTACACGGACA**BHQ2**-3’<br><br>5’**CY5**TGTCCGTGTACACCTGCACCAAACGTGTCCTCGCGTCCGCGTTTTTTTTTTTTTTTTTTTT3’ |
| dUdG-MM | 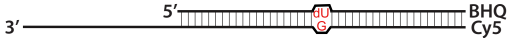 | 819259.2 | 5’GCCCTGCTGCCGACCAACGA**dU**TGGTTACATTCCCGCTGCTG**BHQ2**-3’<br><br>5’***CY5***CAGCAGCGGGAATGTAACCAGTCGTTGGTCGGCAGCAGGGCTTTTTTTTTTTTTTTTTTTT3’ |

**Supplemental Table 4: Individual oligos used to construct longer substrates**

(see Table 1 and 2 above)

| Name | Oligo Sequences |
| --- | --- |
| Duplex 1 | 5’GCCCTGCTGCCGACCAACGATTGGTTACATTCCCGC<br>TGCTGGAGAAACGGCCGAATACA-3’<br><br>5’***p***TCCATGTATTCGGCCGTTTCTCCAGCAGCGGGAATG<br>TAACCAATCGTTGGTCGGCAGCAGGGCTTTTTTTTTTTTT<br>TTTTTTT3’ |
| Duplex 1 PEG | 5’GCCCTGCTGCCGACCAACGA***Sp9***TGGTTACATTCCC<br>GCTGCTGGAGAAACGGCCGAATACA-3’<br><br>5’***p***TCCAAGTATTCGGCCGTTTCTCCAGCAGCGGGAATG<br>TAACCAATCGTTGGTCGGCAGCAGGGCTTTTTTTTTTTTT<br>TTTTTTT3’ |
| Duplex 1 Cy3 | 5’GCCCTGCTGCCGACCAACGA***Cy3***TGGTTACATTCCC<br>GCTGCTGGAGAAACGGCCGAATACA-3’<br><br>5’***p***TCCATGTATTCGGCCGTTTCTCCAGCAGCGGGAATG<br>TAACCAATCGTTGGTCGGCAGCAGGGCTTTTTTTTTTTTT<br>TTTTTTT3’ |
| Duplex 2 | 5’***p***TGGAGAAACGGCCGAATACACC**BHQ2**-3’<br><br>5’***CY5***GGTGTATTCGGCCGTTTC3’ |

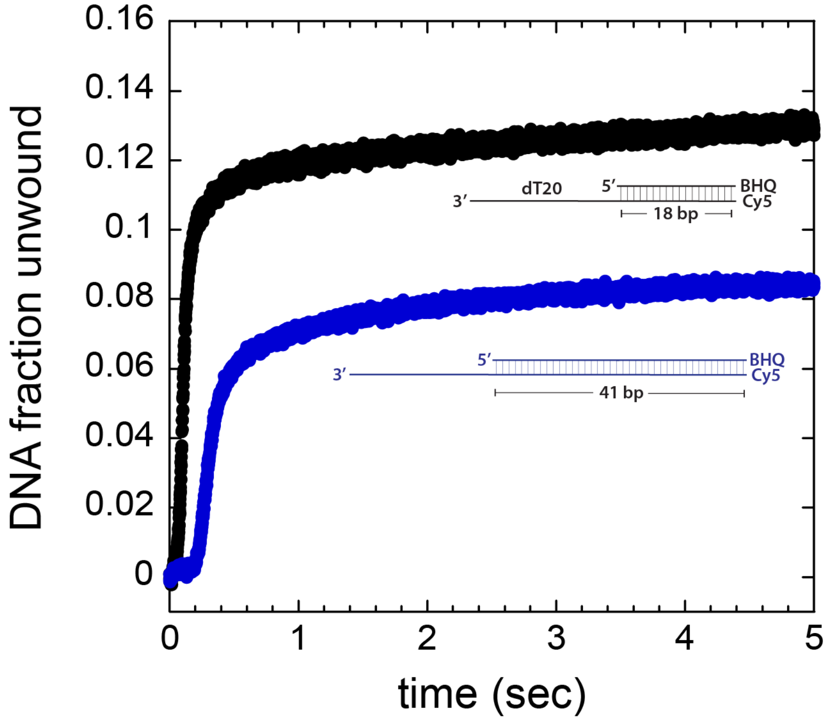


**Supplemental Figure 1: Un-normalized DNA unwinding traces.** Raw fraction DNA unwound as a function of time by *Mtb* UvrD1 is shown on 18 bp (black) and 40 bp (blue) duplexes.

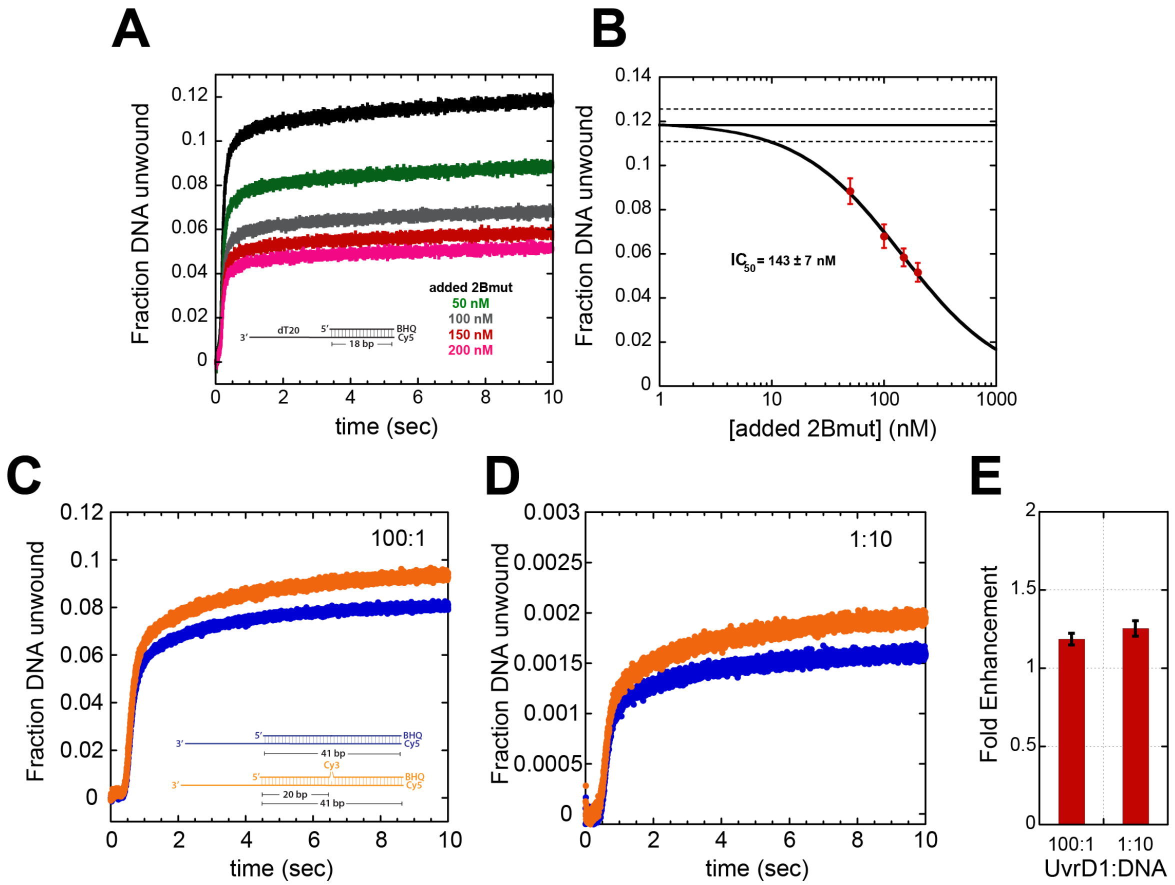


**Supplemental Figure 2: Stimulation of unwinding is an effect on single dimeric enzymes. (A)** Fraction unwound under excess enzyme conditions (black) as a function of the concentration of 2B mutant (C451A) monomeric enzyme as indicated in the legend. **(B)** Quantification of the inhibitory curve from (A). **(C)** Fraction unwound as a function of time in the presence of excess enzyme over DNA substrate (100:1 enzyme:DNA ratio). **(D)** Fraction unwound as a function of time with excess DNA substrate (1:10 enzyme:DNA ratio). **(E)** Quantification of the stimulatory effect under excess enzyme and excess DNA conditions.

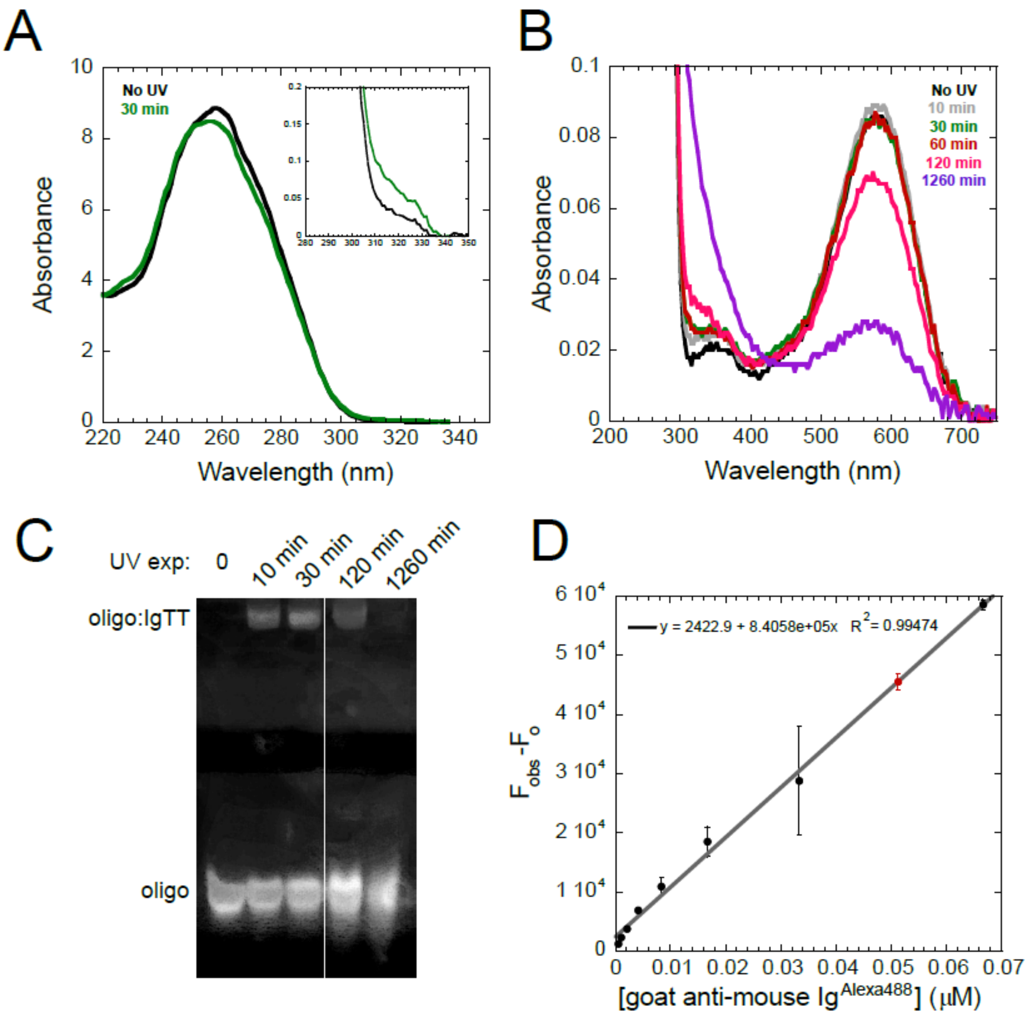


**Supplemental Figure 3: Thymine dimer formation and quantitation. (A)** Absorbance spectrum of oligo for thymine dimer substrate with and without 30 min UV exposure. Inset shows the absorbance change at 325 nm upon UV exposure indicating (6-4) photoproduct formation. **(B)** Absorbance spectrum of oligo over UV-vis range showing change in the black hole quencher 2 absorbance (570nm) upon long UV exposure. **(C)** EMSA of oligo treated with indicated UV exposure bound by antibody (Ig) specific for syn cyclobutane thymine dimer. **(D)** A standard curve measuring the fluoresence of different amounts of Alexa488 labeled antibody is shown in black data points fit by the grey line. The quantitation of our experimentally modified substrate is shown in red.

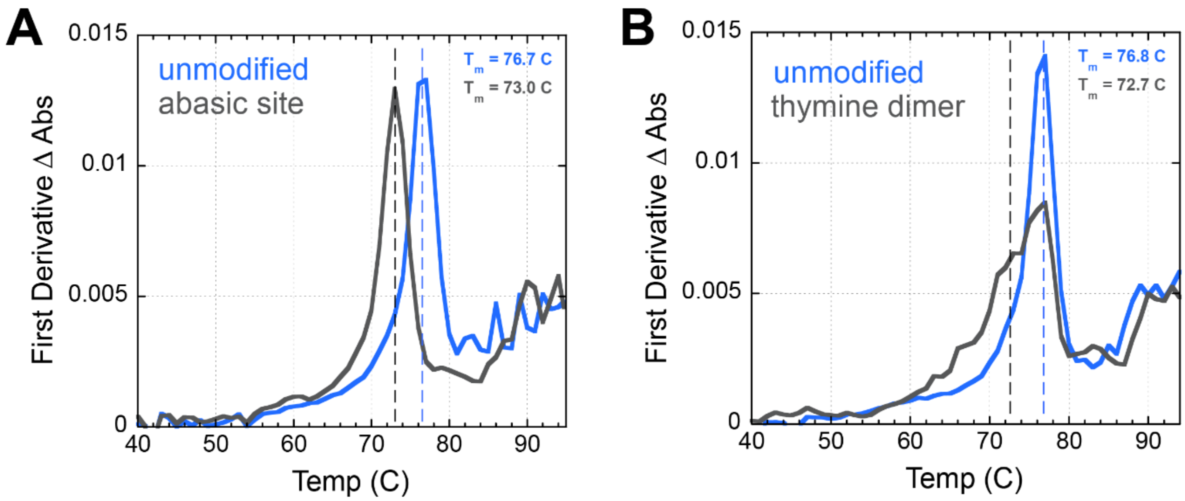


**Supplemental Figure 4: Melting curves comparing the thermodynamic stability of different substrates.** The first derivative of melting curves is plotted as a function of temperature. In each case, the melting temperatures were obtained from maximum(s) in the first derivatives. **(A)** Unmodified (blue) and abasic site (black) DNA substrates. **(B)** Unmodified (blue) and UV irradiated (thymine dimer, black) DNA substrates. Two peaks (modified and unmodified) are observed in the UV irradiated substrate due to incomplete modification.

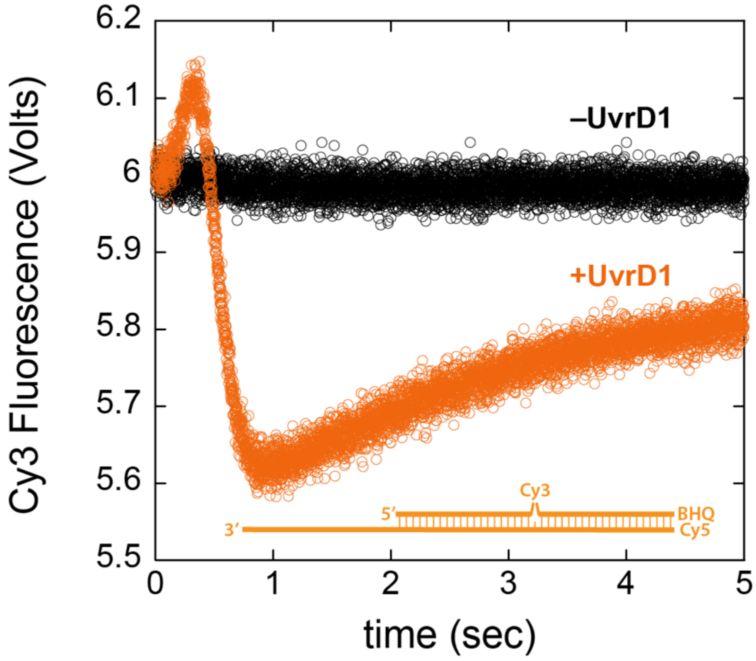


**Supplemental Figure 5: Fluorescent changes are protein dependent.** The fluorescence changes observed during unwinding by UvrD1 (orange) are not present in a control trace without protein (black) on the same substrate.

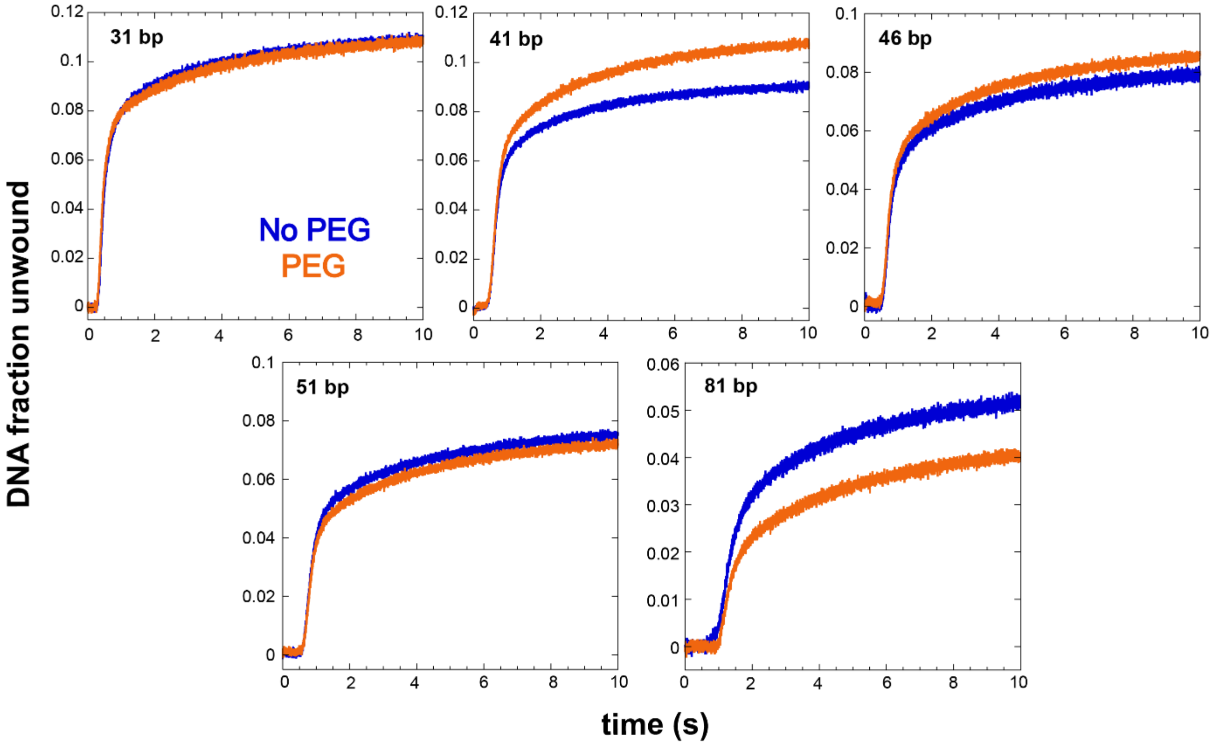


**Supplemental Figure 6: Unwinding curves in the presence and absence of a PEG modification across different lengths.** Examples of traces underlying the data presented in Figure 4 of the main text. The fraction DNA unwound in the presence (orange) or absence (blue) of a PEG3 modification at position 20 are shown for different lengths (indicated in the top left of each plot).

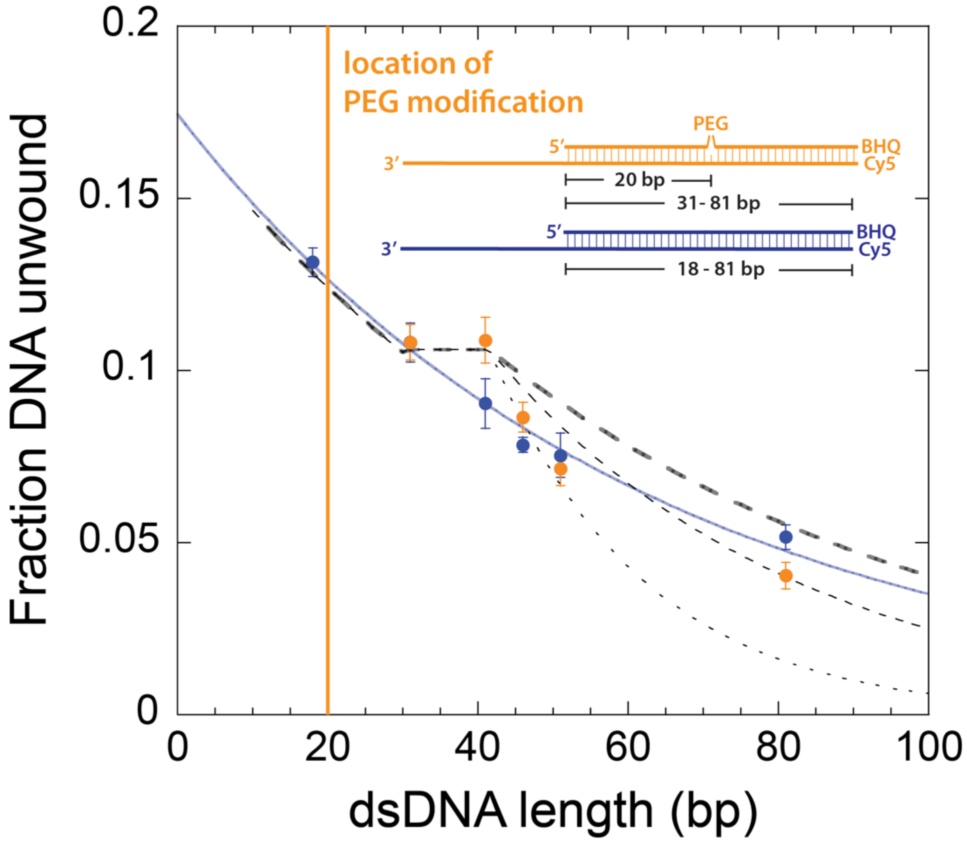


**Supplemental Figure 7: Version of Figure 4 from main text with models of increased dissociation rates 20 bp downstream of the PEG modification.** The longest dashed line (as in Figure 4) represents the case where the dissociation rate returns to the original value that accounts for the basal processivity observed in the absence of modification (blue). The line with the shorter dashes shows the predicted trend when the dissociation constant increases three-fold and the dotted line shows the same, but for a dissociation constant that increases five-fold.

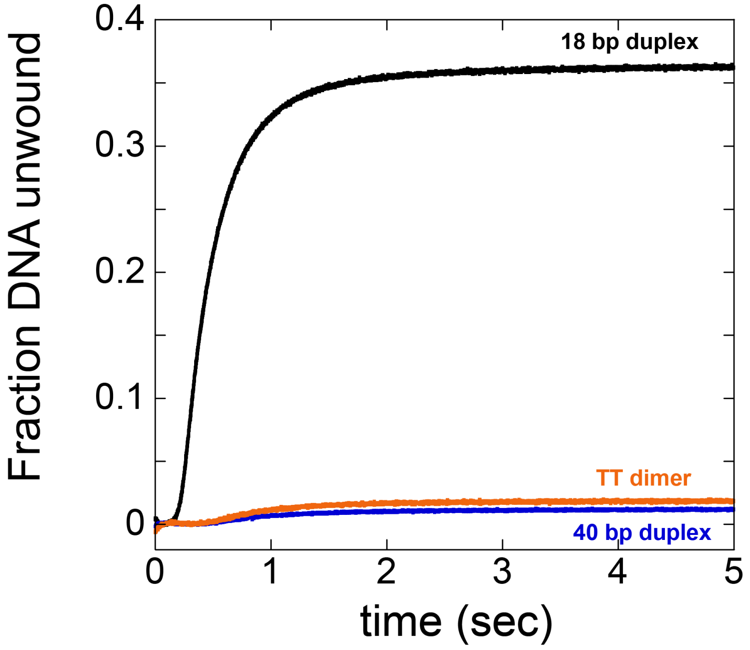


**Supplemental Figure 8: Stimulation of E. coli UvrD unwinding by a thymine dimer in the displaced strand.** Raw fraction DNA unwound as a function of time is shown for *E. coli* UvrD unwinding of an 18 bp duplex (black), a 40 bp duplex (blue) and a 40 bp duplex with a thymine dimer at position 20 (orange). Experiments were performed in 2 mM $MgCl_2$, 1 mM ATP, 1 uM ssDNA trap and 4 μM hairpin trap and the protein and DNA concentrations after mixing were 90 nM and 10 nM respectively.
